# Mutating the arginine residue within the FRNK motif of HC-Pro yield highly attenuated strains that confer complete cross protection against telosma mosaic virus (TelMV) in passion fruit (*Passiflora edulis*)

**DOI:** 10.1101/2024.03.18.585366

**Authors:** Linxi Wang, Wei Shi, Asma Aziz, Xiaoqing Wang, Haobin Liu, Wentao Shen, Aiming Wang, Hongguang Cui, Zhaoji Dai

**Affiliations:** Key Laboratory of Green Prevention and Control of Tropical Plant Diseases and Pests (Ministry of Education), School of Plant Protection, Hainan University, Haikou, Hainan, China; Hainan Key Laboratory for Protection and Utilization of Tropical Bioresources, Institute of Tropical Bioscience and Biotechnology, Sanya Research Institute, Hainan Institute for Tropical Agricultural Resources, Chinese Academy of Tropical Agricultural Sciences, Haikou & Sanya, Hainan, China; London Research and Development Centre, Agriculture and Agri-Food Canada, 1391 Sandford Street, London, Ontario N5V 4T3, Canada

**Keywords:** telosma mosaic virus, TelMV, cross protection, HC-Pro, FRNK, passion fruit, *Passiflora edulis*, mild strain, highly attenuated mutant

## Abstract

Telosma mosaic virus (TelMV, *Potyvirus, Potyviridae*) is an emerging viral pathogen causing a major global threat to passion fruit plantations. However, an efficient strategy for controlling such viruses is not yet available. Cross protection is a phenomenon in which pre-infection of a plant with one virus prevents or delays superinfection with the same or closely related virus. HC-Pro is the potyviral encoded multifunctional protein involved in several steps of viral infection, including multiplication, movement, transmission and RNA silencing suppression. The main hypothesis we tested in this study was whether it is possible to generate attenuated viral strains capable of conferring protection against severe TelMV infection by manipulating the *HC-Pro* gene. By introducing point mutation into the potyviral conserved motif FRNK of HC-Pro, we have successfully obtained three highly attenuated mutants of TelMV (R_181_K, R_181_D and R_181_E, respectively) that can systemically infect passion fruit plants without any noticeable symptoms. Importantly, these mutants confer complete protection against subsequent infection of severe recombinant virus TelMV-GFP, evidenced by no detection of viral RNA or protein of the superinfection virus in the systemic leaves of passion fruit plants in both early and late stages. Lastly, we demonstrated that the HC-Pros harbored by the highly attenuated mutants exhibited reduced RNA silencing suppression activity in *Nicotiana benthamiana* leaves. Altogether, this study provides the first demonstration of the generation of highly attenuated strains for TelMV and highlights key amino acid residue involved in complete cross protection against TelMV, opening a new avenue to fight TelMV in the field.

## Introduction

Cross protection is a phenomenon in which pre-infection of a plant with one virus prevents superinfection (subsequent challenge) with the same or closely related virus. This natural phenomenon was first discovered by Mckinney in 1929, where he found that infection of tobacco plants with one genotype of tobacco mosaic virus (TMV) can protect against the secondary infection with another genotype of TMV (McKinney 1929; Ziebell and Carr 2010). Since then, cross protection has been demonstrated for various plant viruses, animal viruses, bacteriophages and viroids (Folimonova 2013; Ziebell and Carr 2010). Plant virologists have observed this phenomenon in plant viruses belonging to different taxonomic groups, such as TMV in the genus *Tobamovirus*, pepino mosaic virus (PepMV) in the family *Flexiviridae*, citrus tristeza virus (CTV) in the family *Closteroviridae*, papaya ringspot virus (PRSV) and zucchini yellow mosaic virus (ZYMV) in the genus *Potyvirus* (Ziebell and Carr 2010; Folimonova 2013; Chewachong et al. 2015). In the past decades, the use of a mild strain virus as the protective virus has been proven to be practical for cross protection both in the lab and field experiments. There are typical two methods for generating mild strain and/or attenuated strain: 1) searching for naturally existed mild strains, and 2) mutagenesis by physical and chemical means, including UV irradiation, nitrous acid and low/high temperature treatment as well as genetic manipulation of the virus genome (Goh et al. 2023; Raja et al. 2022; Tran et al. 2023).

Passion fruit (*Passiflora edulis*), a perennial evergreen vine growing mainly in tropical and subtropical regions, gains popularity in agriculture and food industries due to its nutritional, medicinal and ornamental values. However, viral diseases are the major constraint for passion fruit production worldwide, including in Brazil, Uganda, China, Vietnam, Thailand and Japan. (Yang et al. 2018; Do et al. 2021; Ochwo-Ssemakula et al. 2012). The most frequent cause of viral diseases in passion fruit belongs to the genus *Potyvirus*, including passion fruit woodiness virus (PWV), cowpea aphid-borne mosaic virus (CABMV), East Asian Passiflora virus (EAPV), Passiflora mottle virus (PaMoV), and telosma mosaic virus (TelMV) (Do et al. 2021; Do et al. 2023; Gou et al. 2023). It is worth mentioning that TelMV is an emerging viral pathogen causing a major global threat to passion fruit plantations. In 2008, TelMV was discovered infecting telsoma (*Telosma cordata, Asclepiadaceae*) in Vietnam, and was first reported to infect passion fruit in Thailand in 2014. The infected passion fruit plants exhibited typical viral symptoms including leaf mottle, mosaic and distortion (Chiemsombat et al. 2014). Since then, TelMV has been reported to infect passion fruit in several places in different provinces across China, including Hainan, Taiwan, Guizhou and Fujian, as well as in Vietnam (Yang et al. 2018; Xie et al. 2020; Do et al. 2021; Zhang et al. 2024; Gou et al. 2023). In 2018, Our lab reported the first complete genome sequence of TelMV infecting passion fruit in Hainan, and subsequently obtained the TelMV infectious clone in 2023 (Yang et al. 2018; Gou et al. 2023). Unfortunately, little is known about the viral disease management of TelMV.

TelMV belongs to the genus *Potyvirus* in the family *Potyviriade*. Potyvirus represents the largest group of known plant RNA viruses and comprises many agriculturally and economically important viruses, including soybean mosaic virus (SMV), potato virus Y (PVY), plum pox virus (PPV) and turnip mosaic virus (TuMV) (Cui and Wang 2019; Yang et al. 2021; Dai et al. 2020). The genome of TelMV consists of a positive-sense single-stranded RNA (+ssRNA) genome of 10049 nucleotides, harboring one large open reading frame (ORF) that encodes a polyprotein that can be further processed into ten mature viral proteins by three viral proteases. In addition, a short ORF (pretty interesting *Potyviridae* ORF, PIPO) embedded within the P3 cistron is predicted to be translated by RNA polymerase slippage (Yang et al. 2018; Chung et al. 2008). The helper component proteinase (HC-Pro), a well-known virus-encoded RNA silencing suppressor (RSS), is the second gene product at the amino terminus of potyviral genome. The potyviral HC-Pro has been revealed to be a multifunctional protein that plays roles in aphid transmission, protease activity, viral replication, viral movement and gene silencing suppression (Valli et al. 2018). Schematically, HC-Pro can be divided into three domains: an N-terminal domain essential for aphid transmission; a central region implicated in various functions, including RNA silencing suppression, enhancement of viral particle yield and viral movement, and a C-terminal region harboring the protease activity and as well as the RSS activity. The central region contains the highly conserved FRNK motif that is essential for RSS activity and symptom development (Shiboleth et al. 2007; Valli et al. 2018). Moreover, mutation of the arginine (R) residue within the FRNK motif has been reported for generating attenuated strains that are capable of conferring cross protection against virus infection by severe strains, including TuMV, ZYMV, and SCMV (Kung et al. 2014; Gal-On 2000; Lin et al. 2007; Xu et al. 2020). Nevertheless, mild/attenuated strains for efficient cross protection have not been reported for TelMV.

The aim of this study was to 1) generate mild/attenuated TelMV strain based on the site-directed mutagenesis of HC-Pro, and 2) to investigate its potential for disease management by cross protection. Owing to the wealth of knowledge of mutation of HC-Pro FRNK motif could generate attenuated strain and be applied in cross protection against severe virus strain, we report here mutation of the arginine residue at the position 181 (R_181_) within the FRNK motif of HC-Pro results in highly attenuated strains that could confer complete cross protection against the subsequent challenge of severe TelMV in passion fruit plants. To our knowledge, this is the first study to demonstrate highly attenuated strains of TelMV and their successful application in cross protection against severe TelMV infection in passion fruit. The significance of these results for promoting the disease management of passion fruit and future prospects are discussed.

## Materials and Methods

### Plant material and growth conditions

Tobacco (*Nicotiana benthamiana*) and passion fruit (*Passiflora edulis*) plants were grown in pots and placed in a plant growth chamber with light for 16 h and darkness for 8 h at a relative humidity of 65%. The temperature was set at 24°C and 22°C during the light and dark periods, respectively.

### TelMV infectious clone and virus inoculation

The wild-type TelMV and GFP-tagged TelMV infection clones used in this study were previously reported by our group in 2023 (Gou et al. 2023).

For the inoculation of *N. benthamiana* plants, *Agrobacterium tumefaciens* harboring virus infection clones were infiltrated into the leaves of three-week-old plant using a needle-less syringe. In brief, *A. tumefaciens* strain GV3101 harboring virus infectious clone was grown overnight on a shaker at 200 rpm. The *Agrobacterium* culture was then centrifuged, washed and resuspended in the agroinfiltration buffer (10 mM MgCl2, 10 mM MES, pH 5.7) supplemented with 200 μM acetosyringone. The optical density (OD600) of *Agrobacterium* was then adjusted to 0.5 for agroinfiltration.

For the inoculation of *P. edulis* plants, the sap prepared from virus-infected *N. benthamiana* plants was rub-inoculated on the cotyledons or true leaves of *P. edulis* plants.

### Generation of TelMV HC-Pro mutants

The point mutations of the Arg_181_ in the FRNK motif of HC-Pro (replacing arginine at position 181 with isoleucine, lysine, aspartic acid, glutamic acid, histidine and alanine, respectively) were introduced into HC-Pro cistron by overlapping PCR using Phusion High-Fidelity DNA Polymerase (Thermo Fisher Scientific) and appropriate primers (Table 1). More specifically, primary PCR was carried out to amplify fragment I and fragment II from pPasFru infectious clone (Gou et al. 2023) using primer pairs a/b and c/d, respectively. Next, overlapping PCR was performed to obtain fragment III using primer set a/d, and fragment I and fragment II as the templates. Then, fragment III and pPasFru infectious clones were digested with *Pst*I and *Sal*I, respectively. Lastly, the two digested fragments were ligated by T4 DNA ligase (Thermo Fisher Scientific) to generate recombinant viruses harboring point mutation. Restriction enzymes were purchased from New England Biolabs Inc. All recombinant viral mutants were verified by double digestion and Sanger sequencing.

**Table 1.**
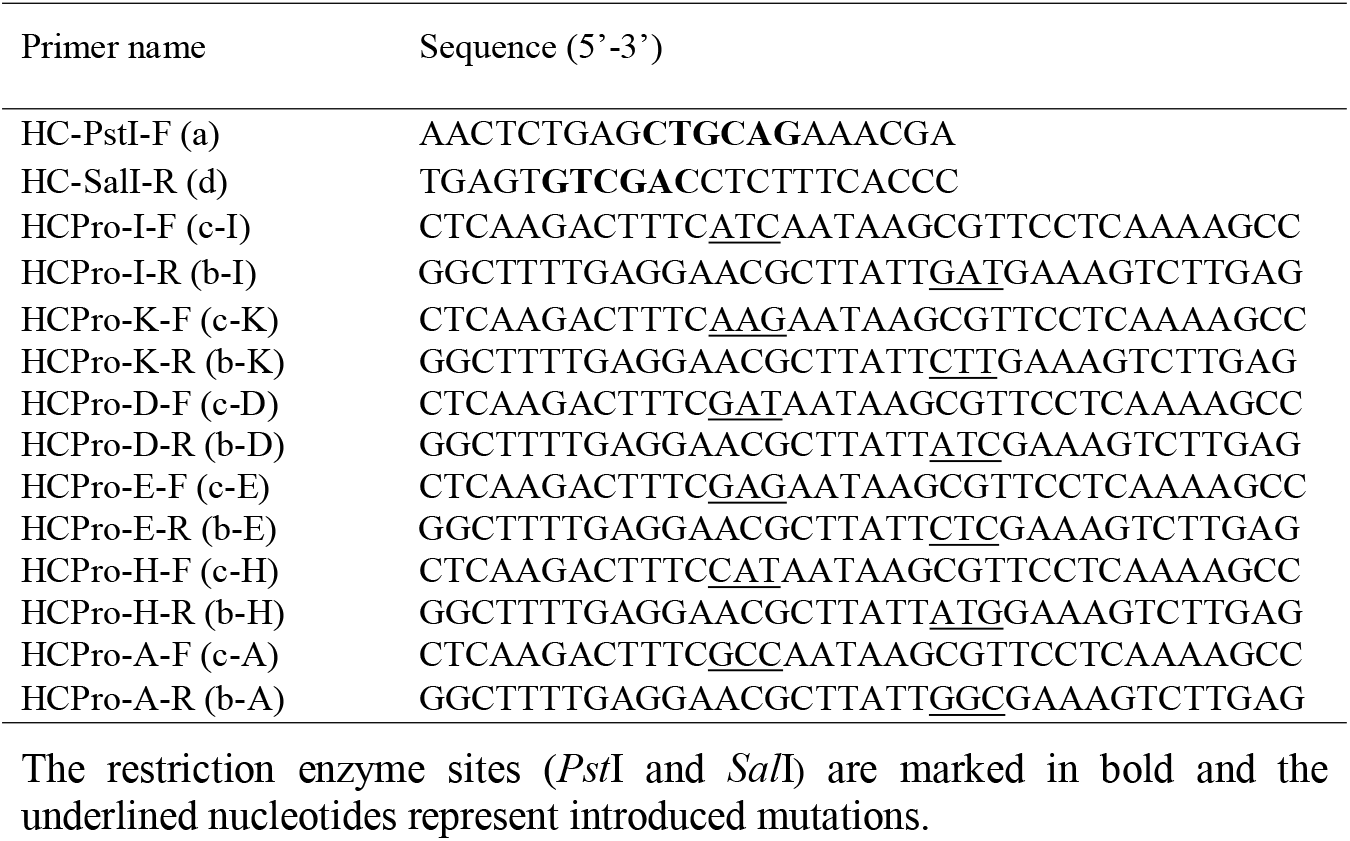
Primers used for the construction of HC-Pro mutants.

### Cross protection assay

For the primary inoculation of *P. edulis* plants, the sap prepared from mutant virus-infected *N. benthamiana* plants was rub-inoculated on the cotyledons of passion fruit plants. To determine whether the passion fruit plants were systemically infected by the mutant virus, systemic leaves were harvested at 12 days post rub-inoculation (dpr) for RNA extraction followed by RT-PCR analysis. Subsequently, the first and second true leaves of these passion fruit plants were further challenge-inoculated with the severe recombinant virus TelMV-GFP (GFP-tagged TelMV infectious clone) (Gou et al. 2023). The cross-protection effect was evaluated by symptom observation under both the regular light and the UV light over an extended period (90 days post challenge). In addition, molecular detection was also carried out to monitor the cross-protection effect. The accumulations of GFP both at the mRNA level and protein level were detected by RT-PCR and Western blot, respectively. The experiment was independently repeated three times, each consisted ten passion fruit plants for the corresponding rub-inoculation.

### Genetic stability assay

The genetic stability of the three highly attenuated mutants (R_181_K, R_181_D, R_181_E) was evaluated by serial passages three times, each with twenty-day intervals. More specifically, *P. edulis* plants were rub-inoculated with the sap prepared from each highly attenuated virus-infected *N. benthamiana* leaves, respectively. Twenty days later, the systemic leaves were collected for grounding and the resulting sap was applied for rub-inoculation on the healthy *P. edulis* plants. The serial passage was carried out three times. Throughout the long passaging period, the inoculated plants remain asymptomatic. Additionally, the systemic leaves are collected from every passage and subjected to RT-PCR analysis, followed by Sanger sequencing to confirm the identity of the mutation.

### RNA silencing suppression assay of HC-Pro mutants

The coding region of both the WT and mutant HC-Pros were amplified from the corresponding virus infection clones by the primer set F1 (5’ATATAGGATCCATGGAACAGAAACTGATCTCTGAAG AAGATCTGTCACTTTCACCTGAGATGCAAT3’)/R1(5’ATATAGGTACCTCAACCAACTCTGTAATGTTTCATTTCTC3’). Note that we introduced the Myc tag-encoding sequence in the primer set. The PCR products were double-digested with *Bam*HI and *Kpn*I, and then inserted into the plant expression binary vector (pCaMterX) (Qin et al. 2020). Each HC-Pro-expression construct was further transferred into *Agrobacterium tumefaciens* GV3101 by the electroporation method.

Next, The WT or mutant HC-Pro was co-expressed with the GFP-expression construct (Li et al. 2014), respectively, by agroinfiltration in *N. benthamiana* leaves. The strong gene silencing suppressor P19 (from tomato bushy stunt virus, TBSV) and empty vector were used as the positive control and negative control, respectively. Lastly, the GFP fluorescence was visualized under UV light starting at 2 dpi and observed over an extended period (11 days). The protein accumulation of GFP was detected by Western blot at 4, 7 and 11 dpi, respectively. The experiment was repeated three times.

### RNA extraction and RT-PCR

Total RNA was extracted from 50 mg leaf tissue of *P. edulis* plants using RNA prep Pure Plant Plus Kit (Tiangen). For first-strand cDNA synthesis, 500 ng of RNA was subjected to reverse transcription reaction using the SuperScript III First-strand Synthesis System (Thermo Fisher Scientific) as instructed. For RT-PCR, primers pairs TelMV-8285-F (5’CCAAAGTTAGAGCCAGAAAGGA3’)/ TelMV-8900-R (5’GAAACCATTCATGACGACATTCA3’) and GFP-F (5’ATGAGTAAAGGAG AAGAACTTTTC3’)/GFP-R (5’TTTGTATAGTTCATCCATGCCATG3’), were used to detect the viral mRNA and GFP mRNA, respectively. A fragment of *EF1* was amplified from passion fruit cDNA using primers EF1-F (5’GGCTGAGCGTGAACGTGGTA3’)/EF1-R (5’CGGCACAATCAGCCTGG GAA3’) and served as the internal control (Zhao et al. 2022).

### Western blot analysis

Western blot analysis was essentially performed as described (Wang et al. 2021).

## Results

### TelMV HC-Pro possesses the highly conserved FRNK motif

First, we investigated whether TelMV HC-Pro contains the FRNK motif by multiple alignments of amino acid sequences of HC-Pro derived from various potyviruses. The alignment results show that the FRNK motif shared over 95% amino acid identity among 143 potyviruses (Fig. 1A). Fig. 1B showed the protein sequence alignment of the partial central domain of HC-Pros harboring the FRNK domain derived from 21 potyviruses. Next, we also conducted the protein sequence alignment analysis on the FRNK motif from all the available TelMV isolates deposited in NCBI databases. We found that all the nine TelMV isolates do contain the FRNK motif and share 100% amino acid identity (Fig. 1C). Taken together, TelMV HC-Pro possesses the highly conserved FRNK motif shared by a large number of potyviruses.

**Fig. 1.**
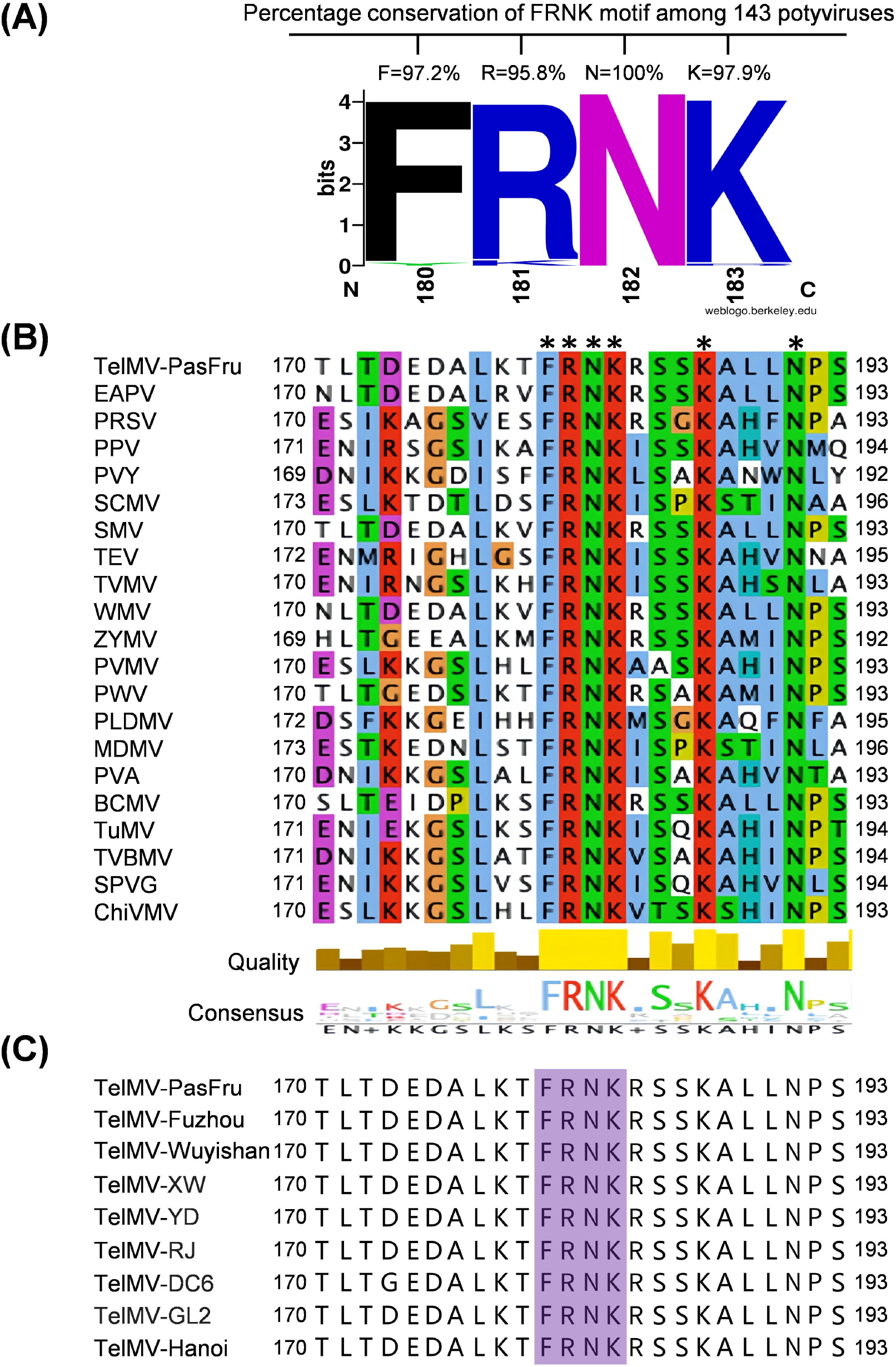
Telosma mosaic virus (TelMV) harbors the highly conserved motif FRNK in potyvrial HC-Pros. A, percentage of amino acid conservation within the FRNK motif of HC-Pros derived from 143 potyviruses. B, protein sequence alignment of the partial central domain of HC-Pros harboring the FRNK domain derived from 21 potyviruses. The abbreviated species names and their GenBank accession numbers are as follows: TelMV (telosma mosaic virus), MG944249; EAPV (East Asian Passiflora virus), AB246773; PRSV (papaya ringspot virus), X67673; PPV (plum pox virus), AY028309; PVY (potato virus Y), X12456; SCMV (sugarcane mosaic virus), AJ297628; SMV (soybean mosaic virus), D00507; TEV (tobacco etch virus), M11458; TVMV (tobacco vein mottling virus), X04083; WMV (watermelon mosaic virus), AY437609; ZYMV (zucchini yellow mosaic virus), AF127929; PVMV (pepper veinal mottle virus), DQ645484; PWV (passion fruit woodiness virus), HQ122652; PLDMV (papaya leaf distortion mosaic virus), JX974555; MDMV (maize dwarf mosaic virus), AJ001691; PVA (potato virus A), AJ296311; BCMV (bean common mosaic virus), AJ312437; TuMV (turnip mosaic virus), AF169561; TVBMV (tobacco vein banding mosaic virus), EF219408; SPVG (sweet potato virus G), JQ824374; ChiVMV (chilli veinal mottle virus), AJ237843. Asterisks indicate identical residues. C, multiple alignments of amino acid sequences of HC-Pros of nine different TelMV isolates, with purple boxes highlighting the FRNK motif identified in the alignments. The GenBank accession numbers are as follows: TelMV-PasFru, MG944249; TelMV-Fuzhou, MK340754; TelMV-Wuyishan, MK340755; TelMV-XW, ON932194; TelMV-YD, ON932196; TelMV-RJ, ON932195; TelMV-DC6, MN316594; TelMV-GL2, MT557572; TelMV-Hanoi, NC_009742.

### Site-directed mutagenesis of TelMV HC-Pro

As the Arginine (Arg, R) residue in the FRNK motif of HC-Pro has been revealed to be an ideal target for generating mild/attenuated strains toward successful cross protection in potyviruses, we hypothesized that the highly conserved Arginine residue could be the ideal candidate for generating mild strain of TelMV in passion fruit plants. To test this hypothesis, we selected the Arginine 181 (R_181_) in the FRNK motif and conducted a comprehensive site-directed mutagenesis in the backbone of TelMV infectious clone (pPasFru) (Gou et al. 2023).

The positively charged R_181_ in the FRNK motif of HC-Pro was mutated into five different residues, including two negatively charged amino acids: aspartic acid (Asp, D) and glutamic acid (Glu, E), the other two positively charged: lysine (Lys, K) and histidine (His, H) and Alanine (Ala, A). In addition, as mutating R to isoleucine (Ile, I), often results in mild symptoms in potyviral infections, we mutated R_181_ to I as well in the current study. In total, six TelMV point mutants (designated R_181_I, R_181_K, R_181_D, R_181_E, R_181_H and R_181_A, respectively) have been successfully created by the site-directed mutagenesis in the background of TelMV infectious cDNA clone (Fig. 2A), supported by the PCR identification (Fig. 2B), double digestion (Fig. 2C), and sequencing validation.

**Fig. 2.**
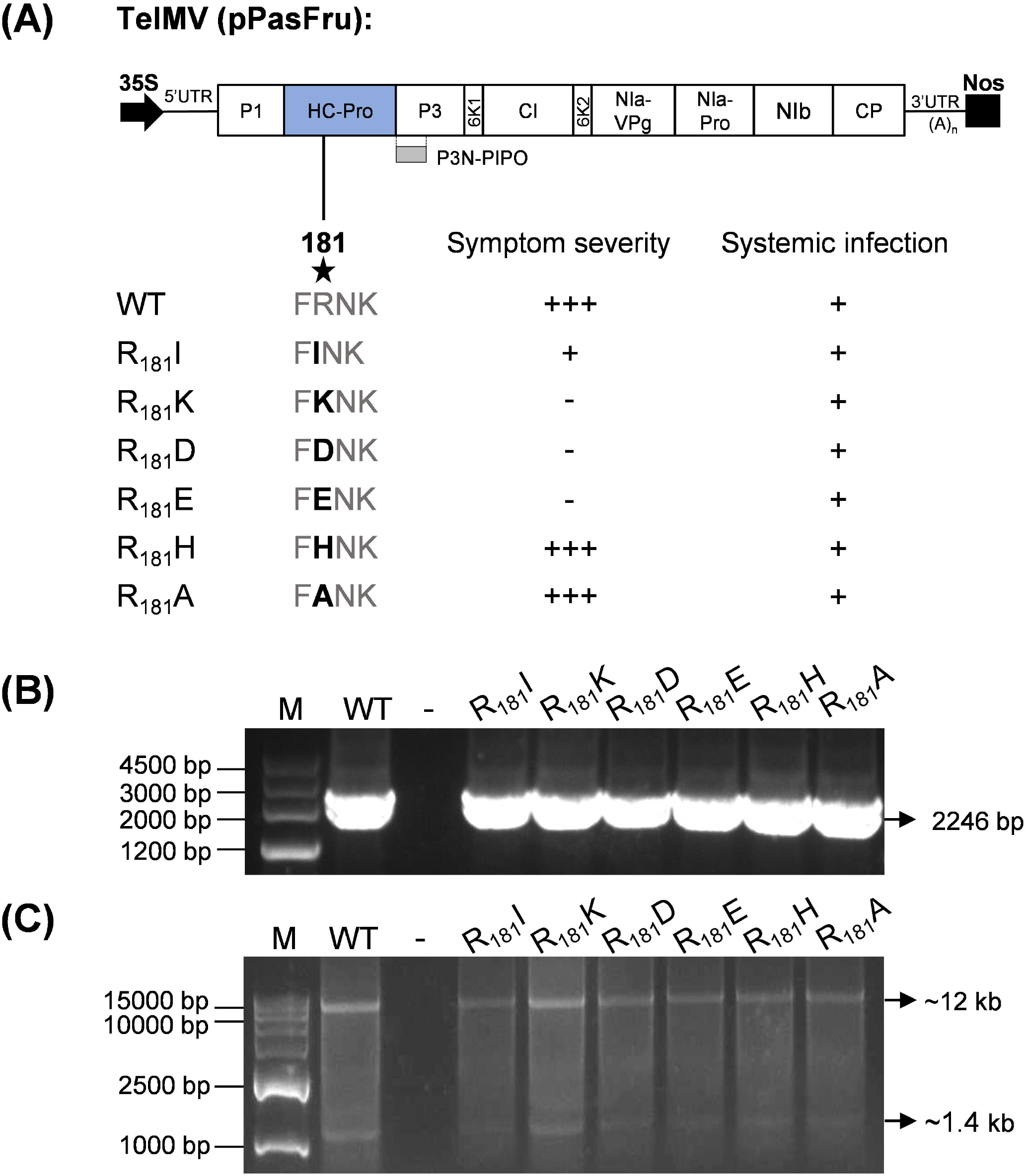
Schematic representation of TelMV HC-Pro point mutants. A, schematic representation of R_181_ mutations in TelMV HC-Pro. The symptom severity is indicated, ranging from severe (+++), moderate (+) to symptomless (–) in passion fruit plants. The systemic infection is determined by the presence (+) or absence (–) of viral RNAs in the systemic leaves of passion fruit plants by RT-PCR analysis. The blue box represents the HC-Pro cistron. 35S: 35S promoter of cauliflower mosaic virus; Nos: nopaline synthase terminator. B, PCR identification of HC-Pro mutants. M: DNA marker. WT: WT virus, serving as the positive control. -: negative control using water as the template. C, double digestion validation of HC-Pro mutants. M: DNA marker. WT: positive control using plasmid of WT virus as the template. -: negative control without any plasmid added.

### Mutation of R_181_ in the FRNK motif of TelMV HC-Pro results in highly attenuated infections in passion fruit plants

To evaluate the infectivity of the six HC-Pro mutants, *Agrobacterium* harboring different mutants were agroinfiltrated into 4-week-old *Nicotiana benthamiana* leaves, respectively. At 5 days post inoculation (dpi), the inoculated leaf samples were ground in pestle and mortar and were subsequently rub-inoculated onto carborundum-dusted leaves of passion fruit plants. Viral infections in passion fruit plants were monitored by visual symptom observation and viral RNA detection by RT-PCR. At 25 days post rub-inoculation (dpr), wild-type (WT) virus-inoculated passion fruit plants exhibited typical viral symptoms exemplified by plant stunting, leaf mosaic and small foliage and distortion (Fig. 3A) when compared to the buffer (Mock)-inoculated control plants. Similar to the WT virus, mutants R_181_H, and R_181_A induced severe symptoms, including plant stunting, leaf mosaic and small foliage. R_181_I caused moderate symptom with inconspicuous leaf mosaic. In contrast, mutants R_181_K, R_181_D and R_181_E induced very mild mosaic symptoms on the systemic leaf during 8 to15 dpr, followed by recovery with no conspicuous symptom in passion fruit plants that are indistinguishable from mock controls at 25 dpr (Fig. 3A), suggesting these mutants are highly attenuated.

**Fig. 3.**
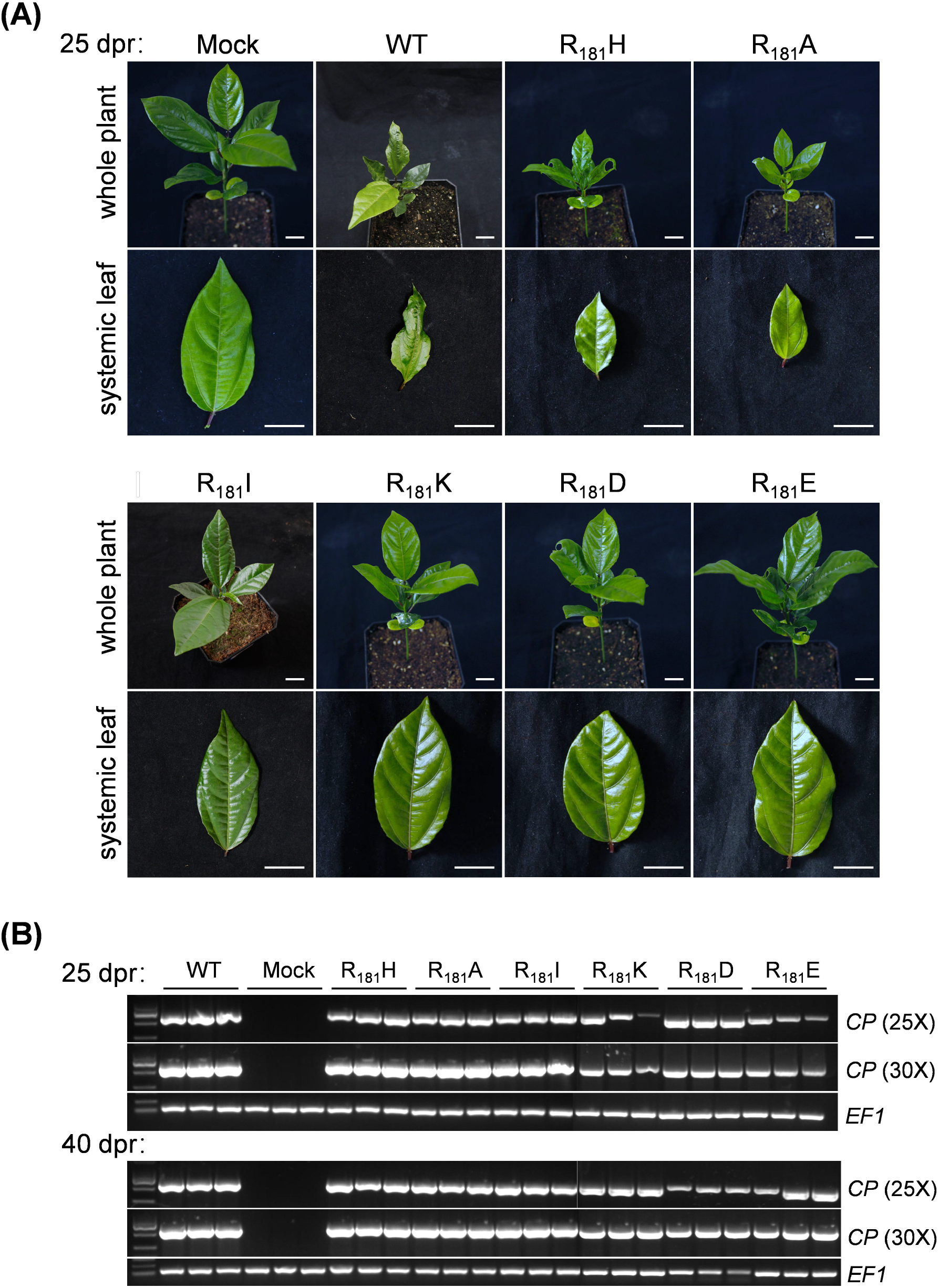
Effect of R_181_ in the FRNK motif of HC-Pro on telosma mosaic virus (TelMV) infection in passion fruit. A, passion fruit plants infected with buffer (Mock), wild-type (WT) virus or HC-Pro mutant virus at 25 days post rub-inoculation (dpr). Scale bar: 2 cm. B, RT-PCR analysis of systemic leaf of HC-Pro mutant virus-inoculated passion fruit plants at 25 dpr and 40 dpr, respectively. WT: wild-type TelMV; Mock: buffer-inoculated plants. A fragment of viral coat protein (CP) gene (687 bp) was used to detect viral RNAs. The *EF1* gene fragment (146 bp) of passion fruit was served as the internal control for RT-PCR analysis.

Intriguingly, RT-PCR analysis of the systemic leaves showed that viral RNA can be detected in the highly attenuated mutants (R_181_K, R_181_D or R_181_E)-inoculated passion fruit plants (Fig. 3B), suggesting these mutants can systemically infect plants without inducing visual symptoms. In addition, at 40 dpr, these mutants did not induce any visual symptoms on passion fruit plants but indeed systemically infected the plants, evidenced by the presence of intensive viral RNA in the newly-emerged leaves by RT-PCR analysis (Fig. 3A, 3B). We constantly monitored the growth of plants till 120 dpr and all plants inoculated by R_181_K, R_181_D or R_181_E were symptomless and indistinguishable from the mock controls.

Taken together, the mutants R_181_K, R_181_D and R_181_E are highly attenuated strains with the capability of systemic infection in passion fruit plants.

### The highly attenuated mutants R_181_K, R_181_D and R_181_E provide complete protection against severe TelMV infection

We next investigated the potential of the highly attenuated strains (R_181_K, R_181_D and R_181_E) to cross protect against severe TelMV infection in passion fruit plants. To this end, ten passion fruit plants were first rub-inoculated with R_181_K, R_181_D or R_181_E, respectively for twelve days to allow for systemic infection (Fig. 4A). Fig. 4B showed the presence of viral RNA in the systemic leaf of highly attenuated mutant-inoculated plants at 12 dpr. Subsequently, passion fruit plants were then challenged with severe recombinant virus TelMV-GFP (Gou et al., 2023) on the upper young leaves and observed for symptom development (Fig. 4A). The experiment was repeated three times with the same results. The unprotected plants developed severe symptoms with plant stunting, mosaic and small foliage (Fig. 4C). In contrast, passion fruit plants pre-inoculated with R_181_K, R_181_D or R_181_E showed no apparent symptom at 30 days post challenge (dpc) or over an extended period (60 dpc) of observation (Fig. 4C). Constantly, no green fluorescence could be observed under UV light in R_181_K, R_181_D or R_181_E-preinoculated plants at 30 dpc and 60 dpc (Fig. 4C), suggesting the highly attenuated mutants R_181_K, R_181_D and R_181_E confer efficient protection against severe TelMV infection in passion fruit plants.

**Fig. 4.**
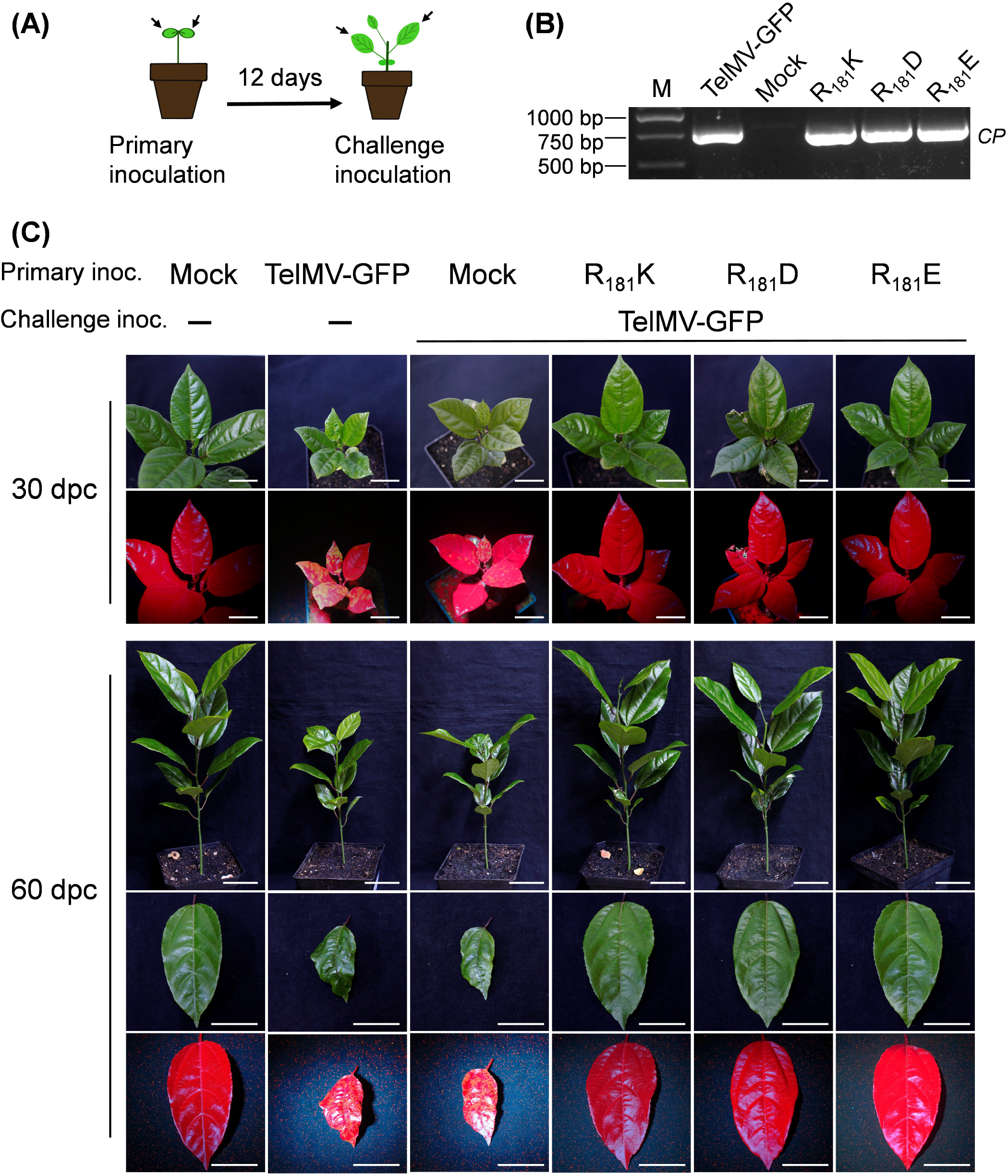
Effect of three HC-Pro mutants on cross protection against severe telosma mosaic virus (TelMV) infection. A, schematic representation of the experimental design for cross protection in passion fruit plants. B, RT-PCR analysis for viral RNA (partial *CP* gene) detection in the systemic leaf of HC-Pro mutant-infected plants before the challenge infection. M: DNA marker. TelMV-GFP: GFP-tagged TelMV infectious clone; Mock: buffer-inoculated plants. C, mutants R_181_K, R_181_D and R_181_E confer efficient cross-protection against TelMV-GFP infection in passion fruit plants at 30 days and 60 days post challenge (dpc), respectively. Scale bar: 3 cm for pictures at 30 dpc and 4 cm for pictures at 60 dpc, respectively.

The visual observation of symptoms was further corroborated with molecular analysis. At 30 dpc and 60 dpc, the RT-PCR results (with 30 cycles) confirmed the presence of GFP gene in Mock plants challenged with TelMV-GFP, but not in R_181_K, R_181_D or R_181_E-preinoculated plants (Fig. 5A), suggesting R_181_K, R_181_D or R_181_E mutants could offer complete cross protection against TelMV-GFP. To further confirm if the viral RNAs of protective virus were present in the systemic leaves of challenged passion fruit plants during the protection period, we used a pair of primer located in the NIb and CP cistron (franking the GFP cistron), respectively for additional PCR analysis. In the non-challenge inoculation sample (TelMV-GFP) and the Mock sample challenged with TelMV-GFP, we detected blight bands corresponding to the size of the NIb-GFP-CP fragment (1431 bp). However, in the R_181_K, R_181_D or R_181_E-preinoculated sample, a smaller band corresponding to the size of NIb-CP fragment (687 bp) was detected (Fig. 5A). These results indicated that the highly attenuated mutants R_181_K, R_181_D and R_181_E can systemically infect the plants with the excellent ability to cross protect severe TelMV infection in passion fruit plants. Furthermore, Western blot analysis also revealed that GFP protein cannot be detected in R_181_K, R_181_D or R_181_E-preinoculated samples (Fig. 5B). Taken together, these results demonstrated that the three highly attenuated strains (R_181_K, R_181_D or R_181_E) could provide complete protection against severe TelMV infection in the passion fruit plants.

**Fig. 5.**
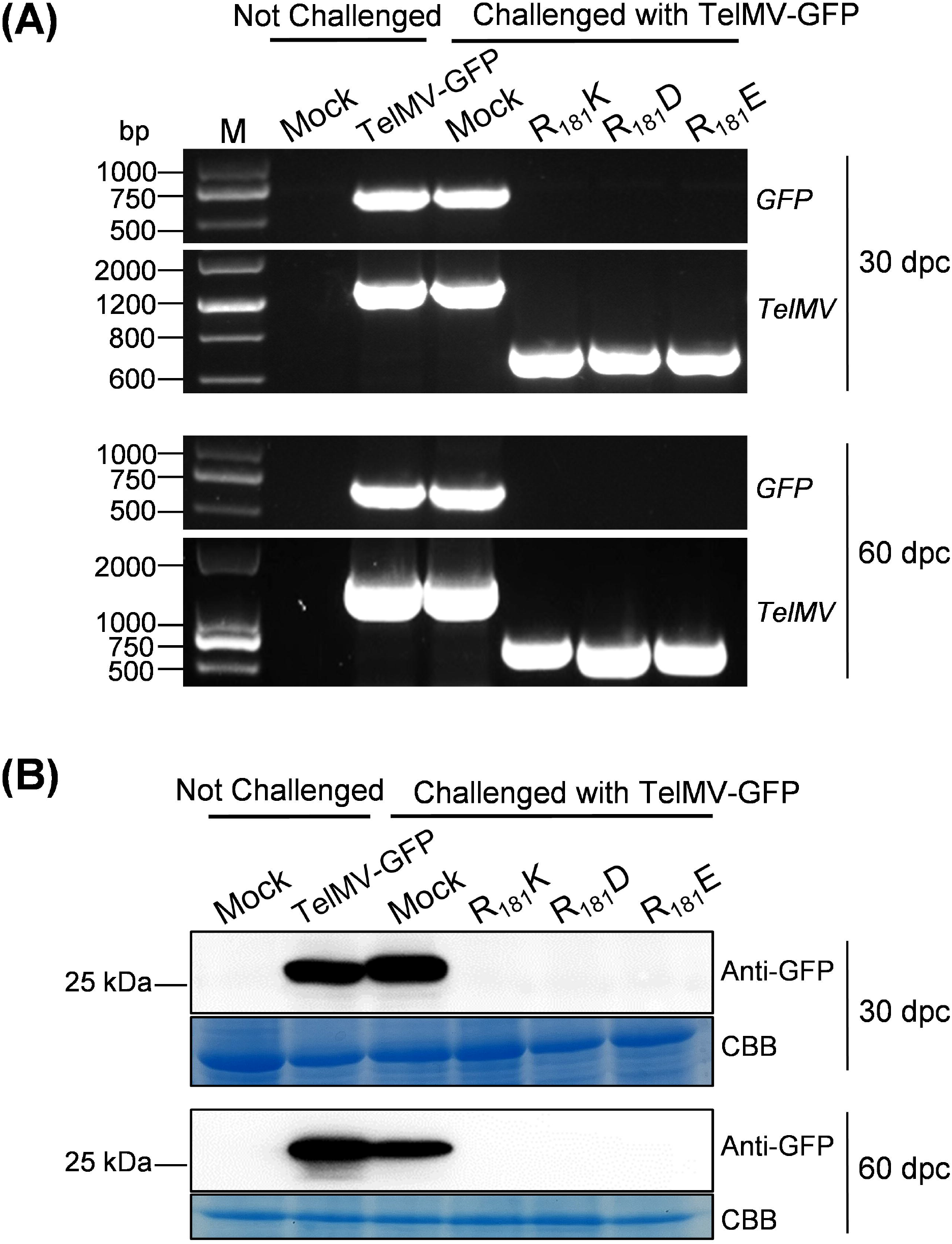
Molecular detection of superinfection in the systemic leaf of passion fruit plant by RT-PCR and Westen blot analyses. A, detection of viral RNA of superinfected passion fruit plants by RT-PCR. B, detection of GFP protein of superinfected passion fruit plants by Western blot. The Coomassie brilliant blue (CBB)-stained Rubisco large subunit (RbcL) serves as a loading control.

### Mutants R_181_K, R_181_D and R_181_E are genetically stable in passion fruit plants

To evaluate whether the highly attenuated strains (R_181_K, R_181_D and R_181_E) were stable and maintained their identity after serial passages in passion fruit plants, the highly attenuated strains were subjected to rub-inoculations in passion fruit plants for three serial passages, each with twenty-day intervals (Fig. 6A). The stability of the highly attenuated strains is determined by both the symptom and molecular analyses. All the plants inoculated with R_181_K, R_181_D or R_181_E were symptomless among all the serial passages tests (Fig. 6B upper panel). Furthermore, the RT-PCR products from passion fruit plants for the serial passages were subjected to sequencing, and the sequence analysis of HC-Pro confirmed that R_181_K, R_181_D and R_181_E remained unchanged during each serial passage (Fig. 6B lower panel). These results revealed that the attenuated strains R_181_K, R_181_D and R_181_E are genetically stable at the genome level through the serial passages.

**Fig. 6.**
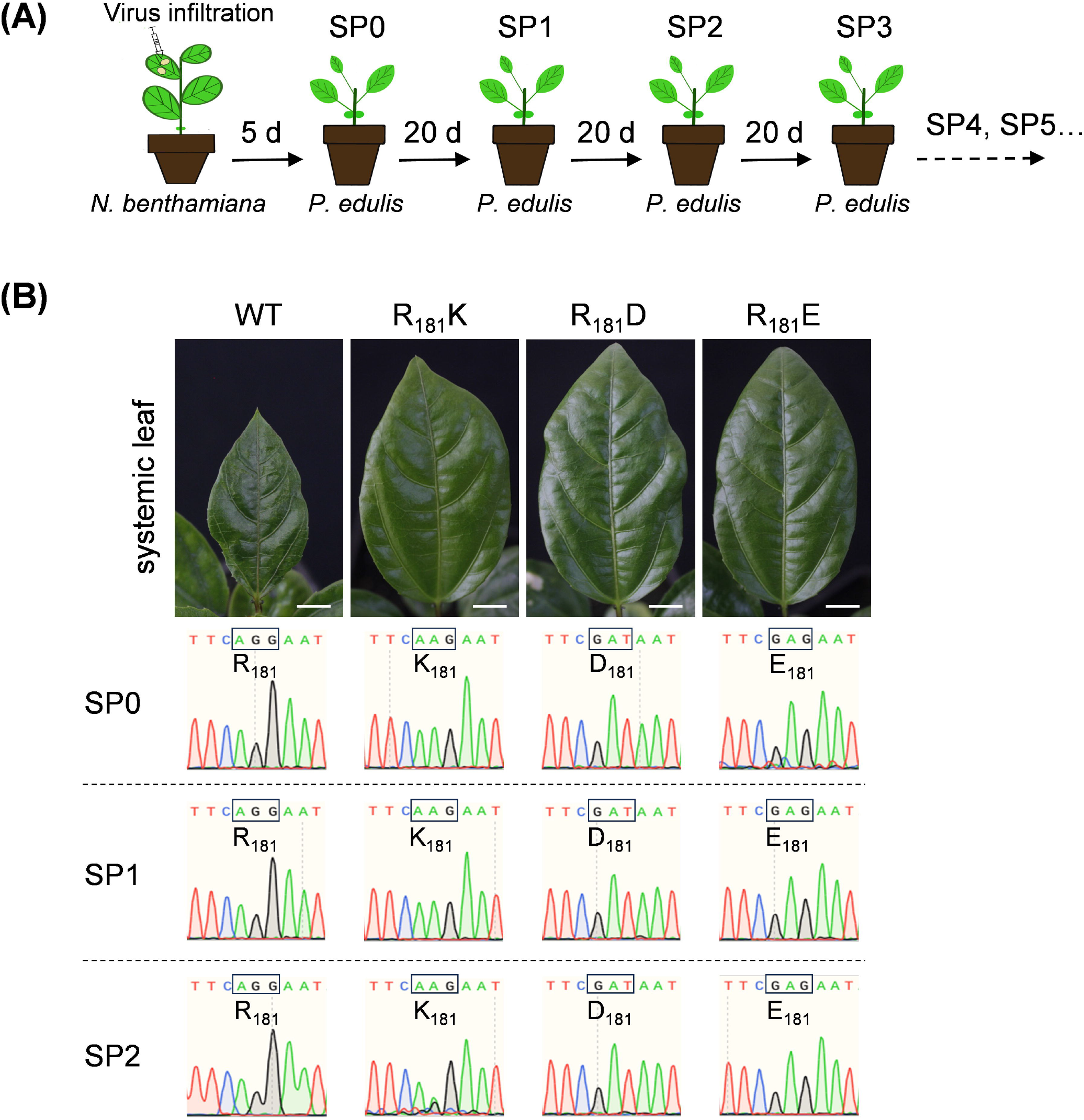
Genetic stability assay. A, schematic representation of genetic stability assay for the highly attenuated TelMV mutants in passion fruit plants. SP: serial passage. B, genetic stability assay by symptom observation and sequence analysis of RT-PCR products of HC-Pro from the highly attenuated mutants-inoculated passion fruit plants. For each serial passage test, the stability of the highly attenuated strains is determined by symptom observation. Scale bar: 1 cm. In addition, the systemic leaves were collected for RNA extraction and the RT-PCR products were subjected to sequencing.

### The Arginine residue within the FRNK motif is critical for RNA silencing suppressor (RSS) activity

As the potyviral HC-Pro is a commonly known RNA silencing suppressor (RSS), we hypothesized that TelMV HC-Pro possesses the RSS activity, and the mutated HC-Pros of the highly attenuated strains execute the eliminated or diminished RSS activity. To this end, the *HC-Pro* gene of WT as well as the mutated viruses were cloned into a Myc-tagged plant expression binary vector (Fig. 7A), respectively and were subsequently examined for RSS activity using a GFP-silencing assay. The Myc-tagged mutant HC-Pro were co-expressed with GFP-expression construct (Qin et al. 2020), respectively, by agroinfiltration in *N. benthamiana* leaves and the GFP fluorescence was visualized under UV light starting at 2 dpi and observed over time for 11 days.

**Fig. 7.**
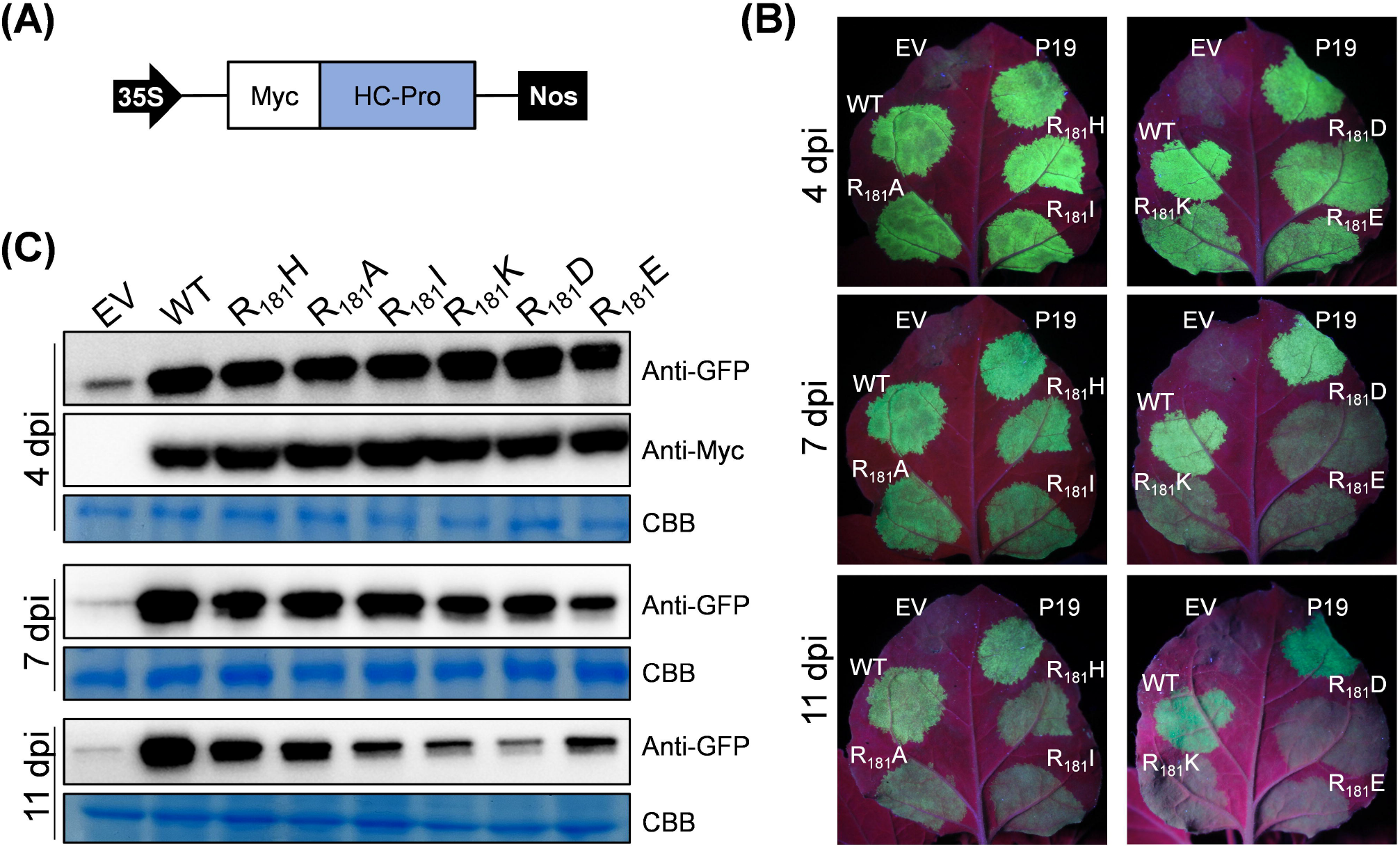
RNA silencing suppression analysis of WT and mutated HC-pros of telosma mosaic virus (TelMV) in plant. A, schematic representation of plant binary vector for HC-Pro expression. B, GFP-silencing assay in *N. benthamiana* leaves. GFP-expression construct was co-expressed with vectors expressing WT HC-Pro or mutated HC-Pros, respectively and monitored under UV light at 4, 7 and 11 days post inoculation (dpi). EV: empty vector; P19: RNA silencing suppressor P19 from tomato bushy stunt virus (TBSV). C, western blot analysis of GFP expression and HC-Pro expression using anti-GFP and anti-Myc at different time points. The Coomassie brilliant blue (CBB)-stained Rubisco large subunit (RbcL) serves as a loading control.

First, by using anti-Myc antibody, western blot analysis showed roughly similar protein levels between the WT and mutant HC-Pros at 4 dpi, suggesting these HC-Pro expression vectors successfully encode the HC-Pro proteins in the GFP-silencing assay experiment (Fig. 7C). Next, GFP fluorescence could be visualized under UV light for both the WT and mutant HC-Pros-expression leaf sample at 4 dpi (Fig. 7B). At 7 dpi and 11dpi, GFP fluorescence in leaf regions co-infiltrated with HC-Pro^WT^ was similar to the fluorescence in regions co-infiltrated with P19 (Fig. 7B), a well-known RSS, indicating TelMV HC-Pro is a very strong RSS.

At 4 dpi, leaf regions co-infiltrated with various mutant HC-Pro exhibited comparable green fluorescence to that with WT HC-Pro, and this is verified by western blot analysis of GFP using GFP antiserum. Interestingly, at 7 dpi, leaf regions co-infiltrated with HC-Pro^R181K^, HC-Pro^R181D^, and HC-Pro^R181E^ exhibited apparent weakened green fluorescence, compared to HC-Pro^WT^. Strikingly, these areas showed hugely diminished fluorescence at 11 dpi (Fig. 7B). In contrast, areas co-infiltrated with the other three mutated HC-pros (HC-Pro^R181H^, HC-Pro^R181A^, and HC-Pro^R181I^) remain the relative strong fluorescence at 7dpi and discernible at 11 dpi (Fig. 7B). This observation is supported by the western blot analysis of GFP at the corresponding time points (Fig. 7C). At 11 dpi, the protein level of GFP from the leaf sample co-infiltrated with HC-Pro^R181K^, HC-Pro^R181D^, and HC-Pro^R181E^ respectively, was apparently lower than that of HC-Pro^WT^ (Fig. 7C).

Taken together, these data indicate point mutants HC-Pro^R181K^, HC-Pro^R181D^, and HC-Pro^R181E^ exhibited weaker RSS function than the WT HC-Pro, demonstrating the residue R_181_ in the FRNK motif is critical for the RSS activity of TelMV HC-Pro.

## Discussion

Telomsa mosaic virus (TelMV) is an emerging viral pathogen that causes devastating economic losses in the passion fruit industry worldwide. However, no TelMV-resistant cultivars are currently available, nor any transgenic approaches (Zhang et al. 2024). Alternatively, cross protection using mild/attenuated strain offers great measures to control virus infection. In the current study, we tested whether mutating HC-Pro could yield mild/attenuated strain for TelMV. We demonstrated for the first time that mutation of Arginine (R) within the highly conserved motif (FRNK) of TelMV HC-Pro could generate highly attenuated strains that elicit full protection against severe TelMV infection in passion fruit plants.

The highly conserved FRNK motif of potyviral HC-Pro is important for viral pathogenicity, mainly evidenced by the previous reports that mutating the Arginine (R) residue resulted in reduced pathogenicity of various potyviruses, including ZYMV, TuMV, PLDMV, EAPV, PaMoV and PRSV (Gal-On 2000; Lin et al. 2007; Shiboleth et al. 2007; Tran et al. 2023; Kung et al. 2014; Tuo et al 2020; Chong et al. 2023; Do et al. 2023). Consistently, in our comprehensive exam on the role of Arginine residue within the FRNK motif of HC-Pro in TelMV infection, three out of six mutants (R_181_K, R_181_D and R_181_E, but not R_181_H, R_181_I and R_181_A) exhibited highly attenuated infections that induced unnoticeable symptom in passion fruit plants (Fig 3A). Intriguingly, mutating R to K in the FRNK motif of ZYMV induced severe symptoms in squash plants (Shiboleth et al. 2007). Similar to our finding, the FKNK mutant of TuMV caused no symptoms in the systemic leaves of *N. benthamiana* plants (Kung et al. 2014), and mutating R to K in the FRNK motif of SCMV induced significantly attenuated infections in maize plants (Xu et al. 2020). It seems the effect of mutating R to K in the FRNK motif on the viral pathogenicity plays a virus-dependent manner. In the case of R to D and R to E mutants, to date, there is only one other report where Kung et al. showed that FDNK and FENK mutants both resulted in the absence of TuMV infection in *N. benthamiana* plants. In our study, the R_181_D and R_181_E indeed induced no noticeable symptom (Fig 3A), but viral RNAs were detected in the systemic leaves (Fig 3B), which further promoted us to test their potential in cross protection against subsequent serve viral infections.

Here, we reported the three highly attenuated mutants (R_181_K, R_181_D and R_181_E) elicited complete cross protection against severe TelMV infection in passion fruit (Fig 4C). It added an example of a single-point mutation of the highly conserved FRNK that could offer full cross protection against severe viral strains. Nevertheless, double mutants with one mutation within the FRNK motif have been used as the mild strains offering cross protection, such as the FKNK/C60A double mutant of SCMV (Xu et al. 2021), R_181_ID_397_N double mutant of PRSV (Tran et al. 2023) and K_53_R_183_EI of papaya leaf distortion mosaic virus (PLDMV) (Tuo et al 2020). However, compared with single-point mutation, the double or triple mutation has the disadvantage of instability that easy to reverse mutate or induce new mutations. Instead, our study revealed three single-point mutants that are stable and could provide full protection against severe virus infection (Fig 4, 5 and 6).

In our study, mutating the Arginine residue (R) to six different amino acid residues resulted in different effects on the viral pathogenicity. The net charge of the HC-Pro mutants is not important for the viral pathogenicity, as mutating the positively charged R_181_ to the negatively charged residues (D and E), and another positively charged residue (K) both resulted in highly attenuated infections in passion fruit plants. As the higher structure of a protein often plays critical roles in protein function in biology, the secondary structure and the 3D structure of both WT and mutant HC-Pros were predicted using the JPred server and I-TASSER server, respectively. Unfortunately, WT HC-Pro and all three highly attenuated mutant HC-Pros (R_181_K, R_181_D and R_181_E) are distinguishable in both the secondary and 3D structures (data not shown). Together, it seems neither the net charge nor the higher structure of HC-Pro is the direct reason for causing highly attenuated infections. Nevertheless, successful viral infections need the assistance of protein-protein interactions among viral proteins as well as the interactions between viral proteins and host proteins. We argue here that it could be the disruption of protein-protein interaction between the HC-Pro and other viral proteins or host proteins is the reason for the highly attenuated pathogenicity of R_181_K, R_181_D and R_181_E.

The potyviral HC-Pro is a well-known virus-encoded RNA silencing suppressor (RSS) (Qin et al. 2020). In the current study, we have revealed the potyviral pathogenicity of TelMV is correlated with the RSS ability of HC-Pro, as the RSS activities of HC-Pros of the three highly attenuated mutants (HC-Pro^R181K^, HC-Pro^R181D^, and HC-Pro^R181E^) are hugely reduced in the GFP-silencing assay (Fig. 7). This was also the case for SCMV where the FKNK mutant of SCMV HC-Pro exhibited very weak RSS activity (Xu et al. 2020). Another study by Kung et al., 2014 also reported that the FKNK mutant of TuMV caused mild symptoms and retained partial RSS function in Arabidopsis. However, it remains unknown whether the RSS activity of the FDNK and FENK mutant of TuMV HC-Pro is hampered (Kung et al. 2014). It seems the FRNK mutant of HC-Pro often exhibited abolished or weak RSS activities. This is supported by the current study of TelMV as well as many other cases among different potyviruses, including WMV, SCMV, TuMV and PaMoV (Xu et al. 2020; Kung et al. 2014; Do et al. 2023). Nevertheless, the FINK mutant of ZYMV HC-Pro maintains RSS activity, despite that the corresponding viral mutant caused attenuation of symptoms in cucurbits (Shiboleth et al. 2007).

In summary, our study presents the first report of generating highly attenuated mutants (R_181_K, R_181_D and R_181_E) for complete cross protection against TelMV in passion fruit plants. Considering the unavailability of efficient strategies for controlling such emerging viral pathogens in passion fruit, it is very important to evaluate the possible application of these highly attenuated mutants in cross protection in the field. Another interesting future direction will be the investigation of the molecular mechanism of virus attenuation. Enormous efforts should be devoted to assessing if any biological aspect of viral infection is disrupted for the attenuated mutants, such as replication and movement, as well as finding host proteins and/or viral proteins sequestered by highly attenuated strains that are critical for subsequent viral infection.

## Acknowledgments

This work was supported by grants from the Hainan Provincial Natural Science Foundation (grant nos. 321QN181 and 322RC564), and the National Natural Science Foundation of China (grant nos. 32360651, 32102157, and 32372484). We thank all the members of the lab for the helpful discussions on this project.

